# Schizophrenia-linked protein tSNARE1 regulates endolysosomal trafficking in cortical neurons

**DOI:** 10.1101/2021.02.09.430442

**Authors:** Melissa Plooster, Martilias S. Farrell, Guendalina Rossi, Hyejung Won, Stephanie L. Gupton, Patrick Brennwald

**Affiliations:** Department of Cell Biology and Physiology, University of North Carolina, Chapel Hill, NC 27599; Department of Genetics, University of North Carolina, Chapel Hill, NC 27599; UNC Neuroscience Center, University of North Carolina, Chapel Hill, NC 27599; Lineberger Cancer Center, University of North Carolina, Chapel Hill, NC 27599

## Abstract

*TSNARE1*, which encodes the protein tSNARE1, is a high-confidence gene candidate for schizophrenia risk, but nothing is known about its cellular or physiological function. We identified the major gene products of *TSNARE1* and their cytoplasmic localization and function in endolysosomal trafficking in murine cortical neurons. We validated four primary isoforms of *TSNARE1* expressed in human brain, all of which encode a syntaxin-like Qa SNARE domain. RNA-sequencing data from adult and fetal human brain suggested that the majority of tSNARE1 lacks a transmembrane domain that is thought to be necessary for membrane fusion. Live-cell imaging demonstrated that brain tSNARE1 isoforms localized to compartments of the endosomal network. The most abundant brain isoform, tSNARE1c, localized most frequently to Rab7^+^ late endosomal compartments. Expression of either tSNARE1b or tSNARE1c, which differ only in their inclusion or exclusion of a Myb-like domain, delayed the trafficking of known endosomal cargo, Neep21, into late endosomal and lysosomal compartments. These data suggest that tSNARE1 regulates endosomal trafficking in cortical neurons, likely through negatively regulating endocytic trafficking or maturation of early endosomes to late endosomes.

## Introduction

Schizophrenia is a severe and heritable neuropsychiatric disorder. Common variation, copy number variation (CNV), and rare loss-of-function (LoF) variation all contribute to its etiology. Genome-wide association studies (GWAS) have identified 145 loci associated with schizophrenia risk, but many of these loci are within non-coding regions; the functional impact of which is not well understood^1^. Functional genomic evidence such as three-dimensional chromatin interactions and quantitative trait loci (QTL) successfully identified 321 high-confidence candidate genes for schizophrenia GWS loci^2,3^. However, the causative role of these genes in schizophrenia pathogenesis remains to be studied, and, for some, even the normal physiological function is not known. The next critical step is to study the function and dysfunction of these high-confidence candidates. *TSNARE1* is one of the high-confidence schizophrenia candidate genes^2^, but its localization and function in the brain are presently unknown.

*TSNARE1* arose from molecular domestication of a Harbinger transposon early in vertebrate evolution^4,5^. Harbinger transposons were an ancient transposon superfamily that encode a transposon protein and a SANT/myb/trihelix DNA-binding protein. Active copies have not been identified in humans, however, some genes in the human genome trace their origin to Harbinger transposons^6^. For example, *TSNARE1* and *NAIF* (nuclear apoptosis inducing factor) contain a domain descended from the DNA-binding protein encoding gene from Harbinger transposons^4,5^. *TSNARE1* encodes a fusion protein of this Myb-like tri-helix domain and a SNARE (soluble N-ethylmaleimide-sensitive attachment receptor) domain.

SNARE proteins are involved in the fusion of opposing lipid bilayers and are functionally conserved across all eukaryotes^7–12^. SNARE proteins promote membrane fusion by assembling into a four-helical SNARE complex. Distinct SNAREs in the complex impart specificity to the membranes that they fuse ^13^. A highly conserved ionic layer within the hydrophobic center of the four-helical bundle of the SNARE complex is critical for SNARE complex assembly and subsequent membrane fusion^14^. The residue that the SNARE protein contributes to this ionic layer distinguishes SNARE proteins as either Q-SNAREs (glutamine) or R-SNAREs (arginine)^15^. A SNARE complex contains one R-SNARE and three Q-SNAREs, further classified as Qa, Qb, and Qc, based on sequence homology. A transmembrane (TM) domain or a membrane attachment site of the Qa SNARE is thought to be critical for allowing the free energy of the SNARE complex formation to be translated into membrane fusion^16^.

Here we investigated the cellular function of tSNARE1. We validated four major gene products of *TSNARE1* in human brain. All isoforms contained a syntaxin-like Qa SNARE domain, and the most abundant transcripts lacked a TM domain, as well as any other predicted site for membrane attachment. We defined the localization and function of tSNARE1 isoforms in the endolysosomal system of cortical neurons, which were consistent with a role for tSNARE1 in negatively regulating early endosome to late endosome membrane trafficking or maturation.

## Results

### *TSNARE1* is a high-confidence candidate gene for schizophrenia risk

Genome-wide association studies (GWAS) of schizophrenia identified 145 loci associated with schizophrenia risk^1^. One of the genome-wide significant (GWS) loci mapped to an intronic region within the gene *TSNARE1*. The association of *TSNARE1* with schizophrenia was supported by multiple lines of functional genomic evidence, including expression quantitative trait loci (eQTL) and three-dimensional chromatin interactions. For example, eQTLs of *TSNARE1* in the adult prefrontal cortex (PFC) colocalized with schizophrenia GWAS (Fig. 1, posterior probability of colocalization [H4] = 0.97)^2^. Transcriptome-wide association studies (TWAS) suggested overexpression of tSNARE1 to be associated with schizophrenia (Bonferroni P = 3.65 × 10^−7^), further confirming the co-localization results^17^. Isoform QTLs (isoQTL) for *TSNARE1* were not associated with schizophrenia, which suggests that it is overexpression of every transcript of *TSNARE1* associated with schizophrenia, rather than overexpression of a specific isoform. Hi-C results from the adult PFC revealed physical interaction between the schizophrenia GWS SNPs and the promoter region of *TSNARE1*, which provides a potential mechanism by which schizophrenia-associated genetic risk factors are linked to *TSNARE1* regulation (Fig. 1)^2^. Therefore, these results collectively suggest that *TSNARE1* is a significant and high-confidence candidate gene for schizophrenia risk. Yet, nothing is known regarding its cellular and physiological localization and function.

**Fig. 1:**
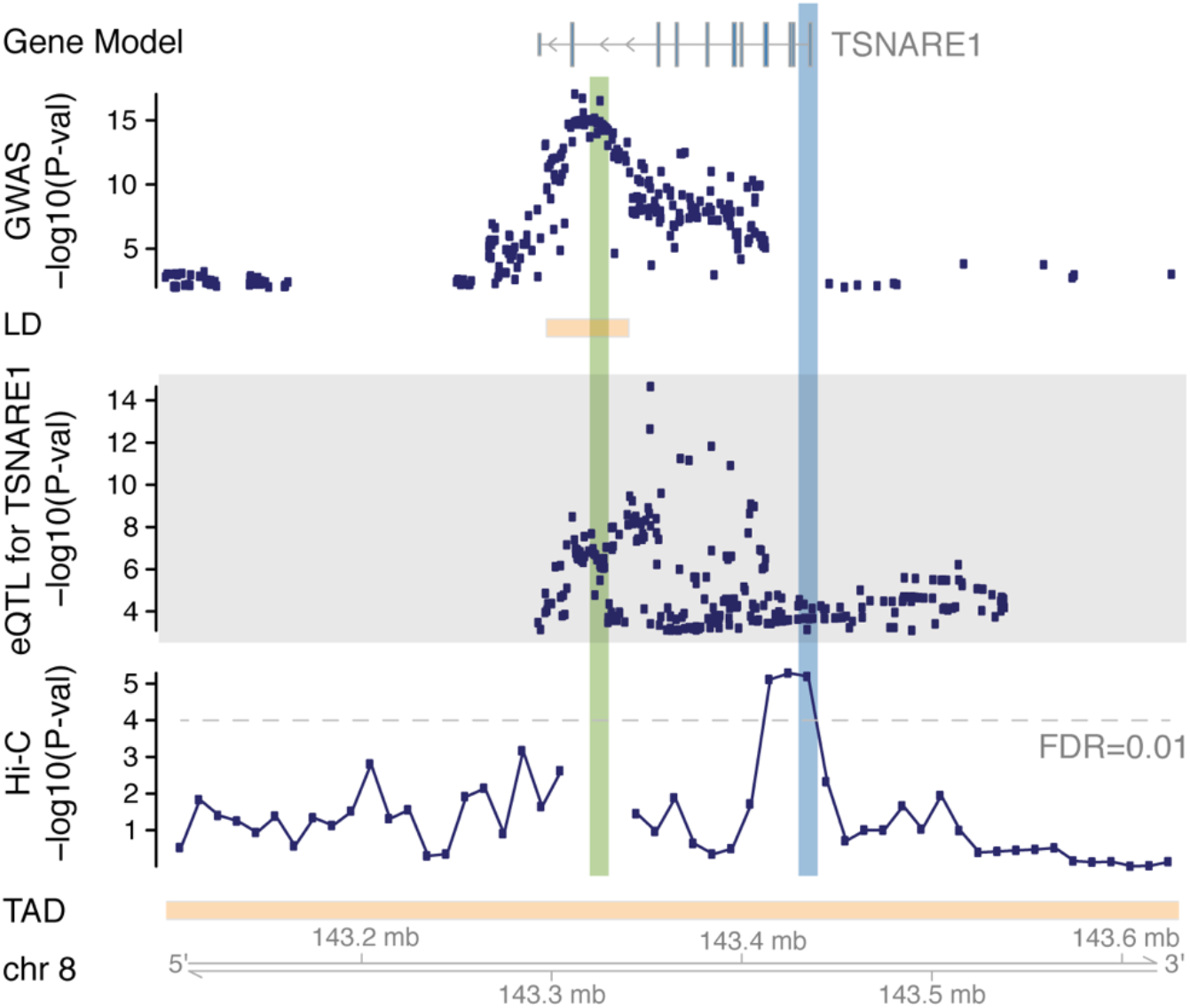
*TSNARE1* is a high-confidence candidate gene for schizophrenia risk. The association between *TSNARE1* and schizophrenia is supported by eQTL and Hi-C data. Hi-C −log10(P-val) denotes the significance of interaction between schizophrenia GWS locus (highlighted in green) and each neighboring bin. *TSNARE1* promoter region is highlighted in blue. GWAS, genome-wide association study; LD, linkage disequilibrium; eQTL, expression quantitative trait loci; TAD, topologically associating domain.

### *TSNARE1* gene expression is enriched in the cortex and in neurons

Because schizophrenia is a neuropsychiatric disorder, we wondered where within the human brain *TSNARE1* is enriched. RNA-sequencing data from the Genotype-Tissue Expression (GTEx) database suggested that *TSNARE1* expression was highest in the cortex as well as the cerebellum (Fig. 2a). We further examined fetal and adult human brain microarrays from the Allen Brain Atlas^18^. Data from three microarray probes are available for *TSNARE1*, but two of the three are within exon 14, an exon predicted to be alternatively spliced that would therefore not reflect expression of all isoforms. However, one probe (CUST_1054_P1416573500) maps to a region within exon 5, which is included in all predicted *TSNARE1* isoforms. This probe revealed enrichment of *TSNARE1* within regions of the cortex in both the fetal and adult human brain (Fig. 2b, Supplementary Fig. 1a-b). We then leveraged single-cell RNA-sequencing results from the human cortex^2^ to investigate cell-type specific expression profiles of *TSNARE1*. We found that *TSNARE1* was highly expressed in neurons and endothelial cells compared with other glial cell types (Fig. 2c). Specifically, *TSNARE1* was more enriched in excitatory neurons than in inhibitory neurons (Fig. 2d).

**Fig. 2:**
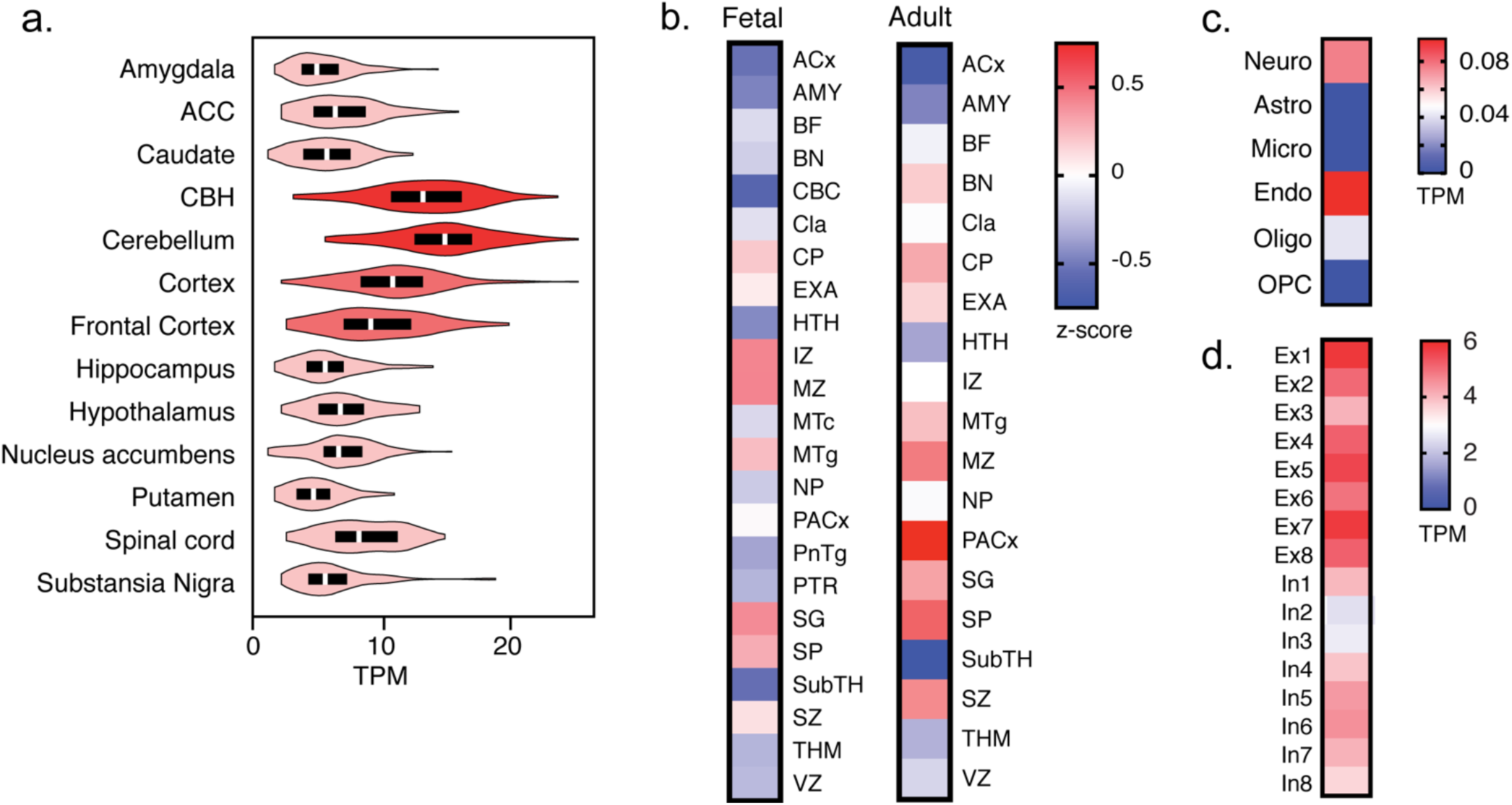
*TSNARE1* gene expression is enriched in the cortex and in neurons. (a) GTEx brain RNA-seq data as plotted for transcripts per million bases (TPM) for *TSNARE1*. ACC, anterior cingulate cortex; CBH, cerebellar hemisphere. (b) Heat map of mean *TSNARE1* expression in brain structures in both the fetal (left) and adult (right) human brain. ACx, allocortex; AMY, amygdaloid complex; BF, basal forebrain; BN, basal nuclei; CBC, cerebellar cortex; Cla, claustrum; CP, cortical plate; EXA, extended amygdala; HTH, hypothalamus; IZ, intermediate zone; MTc, midbrain tectum; MTg, midbrain tegmentum; MZ, marginal zone; NP, neural plate; PACx, periallocortex; PnTg, pontine tegmentum; PTR, pretectal region; SG, subpial granular zone; SP, subplate zone; SubTH, subthalamus; SZ, subventricular zone; THM, thalamus; VZ, ventricular zone. (c) Heat map of mean *TSNARE1* expression in cell types in the brain. Astro, astrocytes; endo, endothelial cells; micro, microglia; neuro, neurons; oligo, oligodendrocytes; OPC, oligodendrocyte progenitor cells. (d) Heat map of *TSNARE1* expression in excitatory and inhibitory neurons. Ex, excitatory neurons; In, inhibitory neurons.

### *TSNARE1* is alternatively spliced, and the majority of transcripts lack a transmembrane domain

Publicly available databases predicted a variety of *TSNARE1* isoforms (RefSeq), but which specific isoforms are expressed in the human brain was not known. The gene *TSNARE1* contains 15 exons (Fig. 3a). Exons 1 and 15 encode the 5’ and 3’ UTR, respectively. Predicted isoforms of *TSNARE1* contained a common N-terminal sequence encoded by exon 2 and one of two C-terminal exons: exon 14 that encodes a transmembrane (TM) domain or exon 13 that encodes a non-TM domain. To identify the isoforms of *TSNARE1* expressed in human brain, we performed PCR using primer pairs against these regions with reverse-transcribed adult cortical and total brain RNA. We validated four isoforms of *TSNARE1* in the human brain that differ in their C-termini and their inclusion or exclusion of exons 3 and 4 (Fig. 3a).

**Fig. 3:**
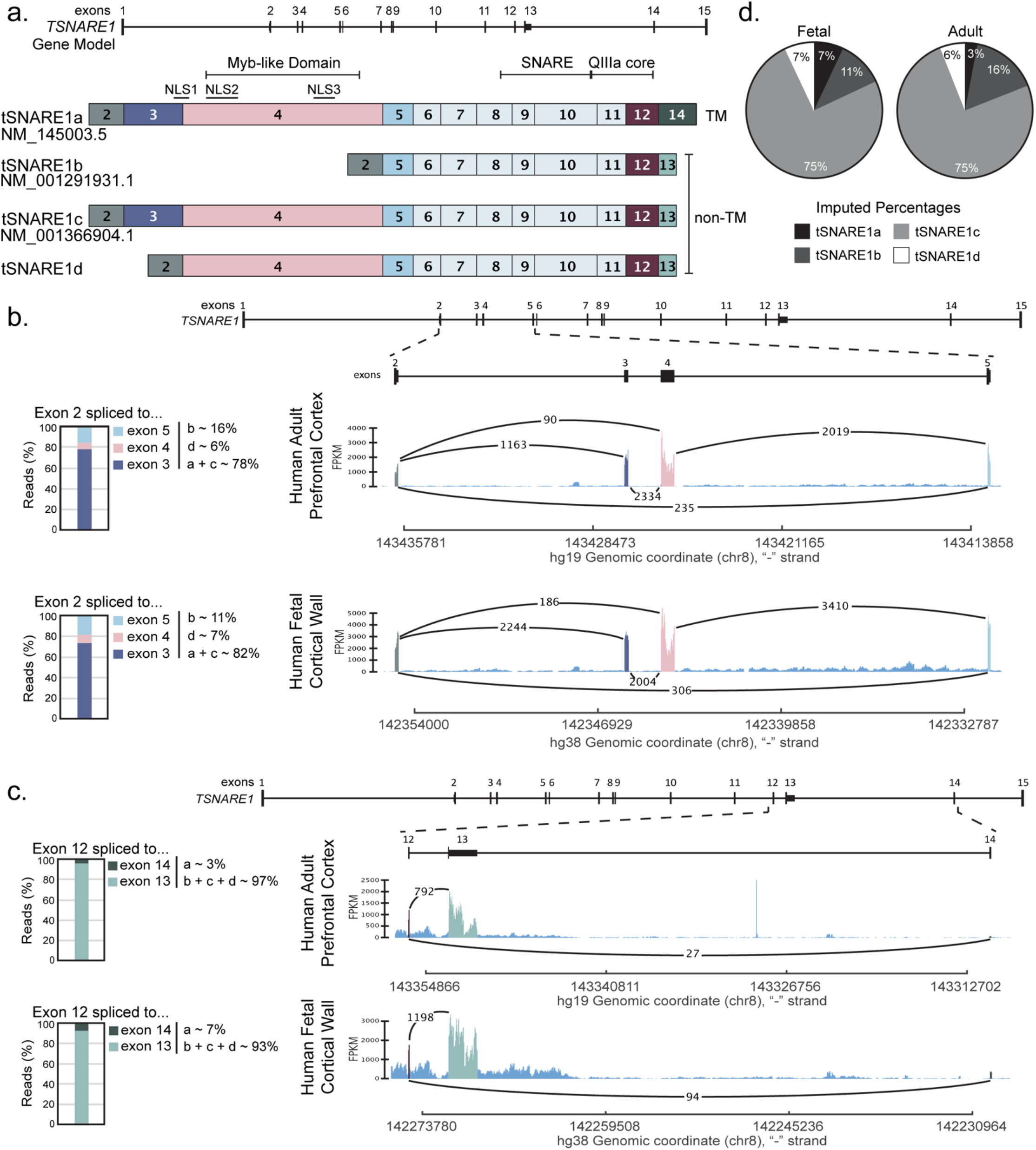
*TSNARE1* is alternatively spliced, and the majority of transcripts lack a transmembrane domain. (a) The gene model for *TSNARE1* and the validated transcripts in adult human brain shown with exon inclusion and the predicted domains and nuclear localization signals (NLS) annotated above. (b&c) Sashimi plot of RNA-sequencing data displaying exon coverage and the number of reads that link two exons for the region from (b) exon 2 to exon 5 or (c) exon 12 to exon 14. (d) The imputed approximate percentages for each isoform calculated by combining the percentages determined in (b) and (c).

To determine the relative abundance of each of these isoforms in the human cortex, we data-mined RNA-sequencing data from the human adult PFC and human fetal cortical wall^2,19^. We examined alternatively spliced regions using sashimi plots showing exon abundance and the number of reads overlapping two exons (Fig. 3b&c). In the region between exons 2 and 5, the majority of reads included both exons 3 and 4, but some skipped both exons or just exon 3 (Fig. 3b). In the other alternatively spliced region between exon 12 to exon 14, the majority of paired reads in exon 12 spliced to exon 13, and the abundance of reads in exon 14 was less than exon 13 (Fig. 3c). These data suggested that the majority of *TSNARE1* mRNA in the adult (~97%) and fetal (~93%) cortex contain the exon encoding a non-TM C-terminal domain. In agreement, the Allen Brain Atlas microarray data demonstrated that the probe common to all isoforms (CUST_1054_P1416573500) had higher expression than either of the two probes that map to the sequence encoding the TM domain in adult brain tissues (Supplementary Fig. 2a). This finding was not limited to the brain. RNA-sequencing data from GTEx revealed that for every tissue, the non-TM encoded by exon 13 had higher expression than the TM domain encoded by exon 14 (Supplementary Fig. 2b) ^20^. From these data, we imputed the approximate percentages of each isoform (Fig. 3d). The most abundant isoform was tSNARE1c, which includes both exons 3, 4, and 13 (~75%) (Fig. 3d).

### *TSNARE1* encodes a SNARE and Myb-like domain

*TSNARE1* encodes two predicted functional domains: a Myb-like domain and a SNARE domain (Fig. 3a). Isoforms that include exon 4 contain a predicted Myb-like domain, and all isoforms of tSNARE1 contain a Qa SNARE domain encoded within exons 8-12, with the core of the SNARE domain encoded by exons 10-12. The observation that the majority of *TSNARE1* mRNA lacked a sequence to encode for a transmembrane domain or any predicted site for membrane attachment is surprising. We wondered whether tSNARE1 would localize as expected for a membrane-bound protein, as discrete puncta in the cytoplasm. Transfection of GFP-tSNARE1 isoforms into murine cortical neurons revealed two distinct populations of tSNARE1 (Fig. 4a). All isoforms exhibited punctate localization within the cytoplasm, primarily within the perinuclear region of the soma. The most abundant brain isoform, tSNARE1c, also localized to the nucleus, consistent with the inclusion of the Myb-like domain.

**Fig. 4:**
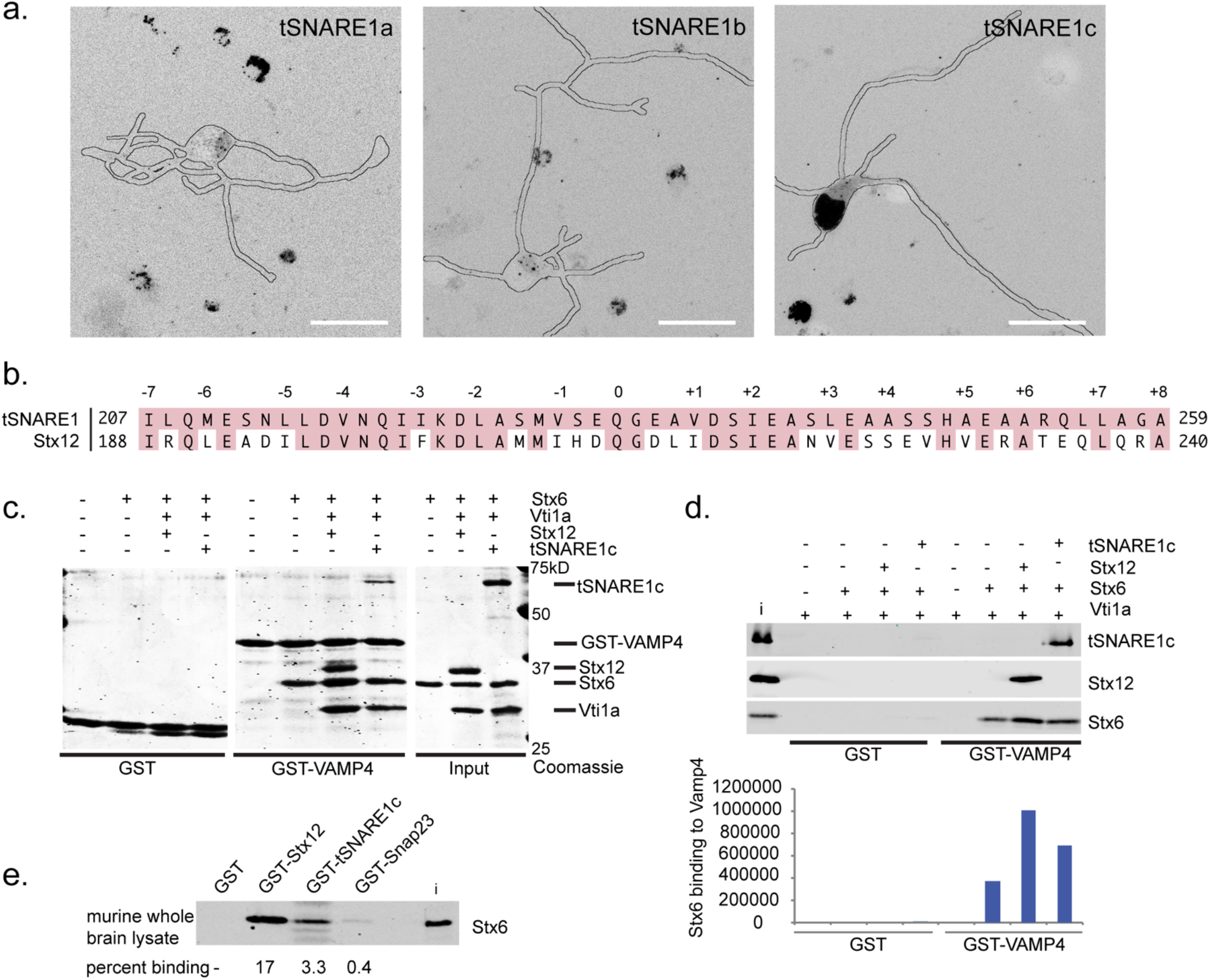
tSNARE1 is competent to form SNARE complexes with Stx6, Vti1a, and VAMP4. (a) Intracellular localization of GFP-tSNARE1 isoforms in murine cortical neurons at 2DIV, with cell outline in black. Scale bar=25μm. (b) Protein alignment of the SNARE domains of tSNARE1 and Stx12. (c) SNARE assembly assays with various recombinant endosomal SNARE proteins. Different combinations of soluble SNARE proteins were mixed with GST-Vamp4 or control GST protein immobilized on beads. Following incubation, beads were washed and assembled complexes visualized by Coomassie staining following SDS-PAGE (see material methods). (d) Assembled SNARE complexes were analyzed by immunoblot analysis using antibodies against specific soluble SNAREs as indicated. Quantitation of the Stx6 incorporation into SNARE complexes is shown below the immunoblot. This allows direct comparison of the ability of tSNARE1c and Stx12 to stimulate incorporation of Stx6 into 4 helical SNARE complexes. (e) Assembly of endogenous Stx6 from mouse brain lysates into heterologous SNARE complexes with recombinant GST-tSNARE1c and GST-Stx12.

We wondered which domain might be more critical for tSNARE1 function. We expect a domain would be less tolerant of variation and have more sequence conservation if it is more critical to the function of tSNARE1. We found that both domains have a positive phyloP score, suggesting both the SNARE domain (Supplementary Fig. 3a, mean=0.449 +/− 0.116 SEM) and the Myb-like domain (Supplementary Fig. 3b, mean=0.338 +/− 0.066 SEM) were evolutionarily conserved. A comparison of copy number variation (CNV) (Supplementary Fig. 3c) and minor allele frequency (Supplementary Fig. 3d) between the SNARE domain and the Myb-like domain suggested that the SNARE domain may be under more evolutionary constraint, but the results were not significant. Since all major products of *TSNARE1* contain the region encoding the SNARE domain, we proceeded to interrogate the role of this domain of tSNARE1 in further detail.

### tSNARE1 is competent to form SNARE complexes with Stx6, Vti1a, and VAMP4

To our knowledge, tSNARE1 is the first Qa SNARE identified that lacks a TM domain or any site for membrane attachment. However, the cytoplasmic localization of tSNARE1 was consistent with the localization of a membrane-bound protein. To confirm that tSNARE1 was competent to form SNARE complexes, we performed biochemical assays with endogenous and recombinant SNARE proteins *in vitro*. The SNARE domain of tSNARE1 shares its closest similarity with syntaxin 12 (Stx12), a QaIII SNARE protein (Fig. 4b). Recombinant Stx12 forms a SNARE complex with recombinant endosomal SNARE proteins Stx6, Vti1a, and GST-VAMP4^21^. Recombinant tSNARE1c similarly formed a SNARE complex with these recombinant endosomal SNARE proteins (Fig. 4c). Recombinant Stx12 potentiated the binding of recombinant Stx6 to GST-VAMP4 *in vitro* (Fig. 4d). Similarly, recombinant tSNARE1c potentiated this binding, although not to the degree of Stx12 (Fig. 4d). Finally, tSNARE1c-GST specifically precipitated the endogenous SNARE protein Stx6 from embryonic murine whole brain lysate (Fig. 4e). These data suggested tSNARE1 was competent to form SNARE complexes with both recombinant and endogenous endosomal SNARE proteins *in vitro*.

### tSNARE1 localizes to markers of the endosomal network

We next sought to determine the intracellular localization of tSNARE1, as proteins typically localize to sites of function. Considering that patients with schizophrenia present cortical abnormalities, excitatory pyramidal cortical neurons are implicated in schizophrenia, and tSNARE1 is enriched in the cortex and in excitatory neurons (Fig. 2), we measured the localization of GFP-tSNARE1 specifically in murine pyramidal-shaped excitatory cortical neurons^2,3,17,22–26^. Because tSNARE1 shares the most sequence similarity with endosomal SNARE protein Stx12 and can interact with Stx12 cognate SNAREs *in vitro*, we hypothesized that tSNARE1 might function within the endolysosomal system. The endolysosomal system is responsible for sorting, recycling, and degradation of internalized cargo. Internalized cargo arrives at Rab5^+^ early endosomes and can be recycled back to the plasma membrane via two separate routes, Rab11^+^ recycling endosomes or Rab4^+^ rapid recycling compartments. Alternatively, cargo is sent from Rab5^+^ early endosomes to Rab7^+^ late endosomes and LAMP1^+^ lysosomes for degradation. To determine to which compartment tSNARE1 localizes, we cotransfected E15.5 murine cortical neurons with GFP-tSNARE1 isoforms and spectrally-distinct red-tagged markers of the endosomal network (Fig. 5a, Supplementary Figure 4a-b).

**Fig. 5:**
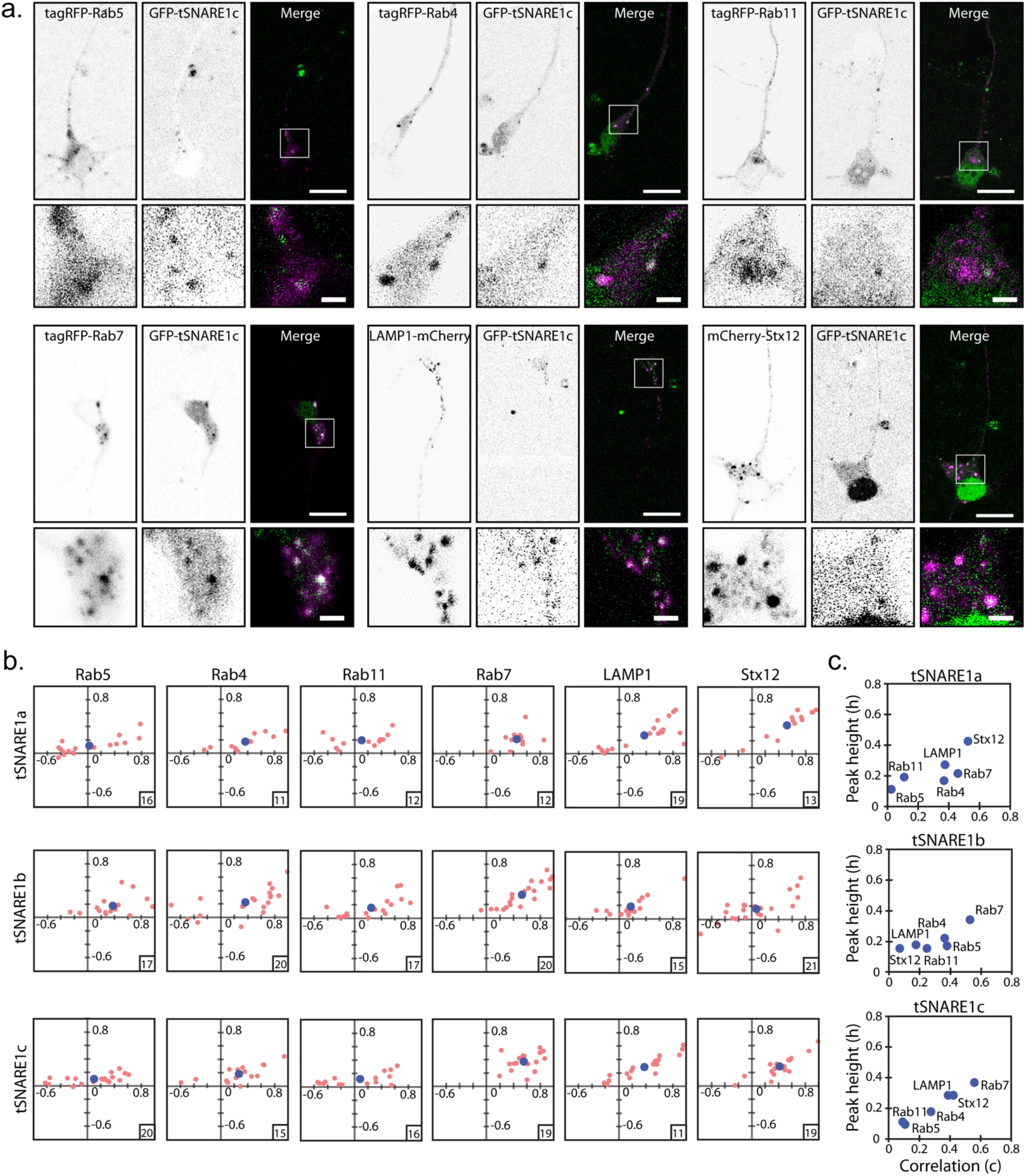
tSNARE1 localizes to markers of the endosomal network. (a) Example images of murine cortical neurons transfected with GFP-tSNARE1c and various red-tagged endosomal markers, with a zoomed in region of the soma marked by a white box. Scale bar=15μm, 2.5μm (zoom). (b) Quantification of colocalization using peak height and correlation of tSNARE1 with each marker. The number of n is boxed on each individual distribution. (d) Averages of individual distributions as shown in (b).

We noted that the majority of tSNARE1 signal localized to the perinuclear region of the soma. We quantified colocalization in the soma using a custom image analysis software that specifically measured colocalization of puncta, which represent membrane-bound organelles^27^. Each isoform of tSNARE1 colocalized positively with a number of the endosomal markers, but the isoforms displayed unique localization patterns (Fig. 5b). tSNARE1b and tSNARE1c, which lack a TM domain, localized most frequently to compartments of the late endosome, marked by Rab7^+^ puncta (Fig. 5c). tSNARE1a, the TM-domain containing isoform, colocalized markedly with Stx12 (Fig. 5c), which itself localized to multiple endosomal compartments. These data indicated that the most abundant isoforms of tSNARE1 in the brain localized to Rab7^+^ late endosomal compartments.

### Overexpression of tSNARE1 delays trafficking of Neep21 into late endosomal and lysosomal compartments

Because tSNARE1b and tSNARE1c most frequently populated compartments of the late endosome, we hypothesized that tSNARE1 regulates trafficking at this compartment. *TSNARE1* is absent from several vertebrate genomes (Ensembl), including mice, perhaps due to a transposition event early in these lineages. We utilized the mouse to exogenously express GFP-tSNARE1c in neurons and compare to a “natural” knockout to interrogate its effect on endosomal trafficking. Because SNARE machinery is conserved across all eukaryotes, we expected tSNARE1 to localize to similar compartments and interact with similar binding partners as it would in human neurons. To determine how tSNARE1 regulates trafficking through the endosomal system, we quantified how overexpression of tSNARE1 altered the trafficking of endosomal cargo protein Neep21. Neep21 is a Type II transmembrane protein, which is rapidly endocytosed and trafficked from the early endosome to the late endosome and the lysosome where it is degraded^28,29^. We measured the localization of a transient population of Neep21-HaloTag, marked by a pulse of a cell-impermeable fluorescent HaloTag ligand, as it trafficked through the endosomal network with or without GFP-tSNARE1 isoforms (Fig. 6a-c, Movies 1-3).

**Fig. 6:**
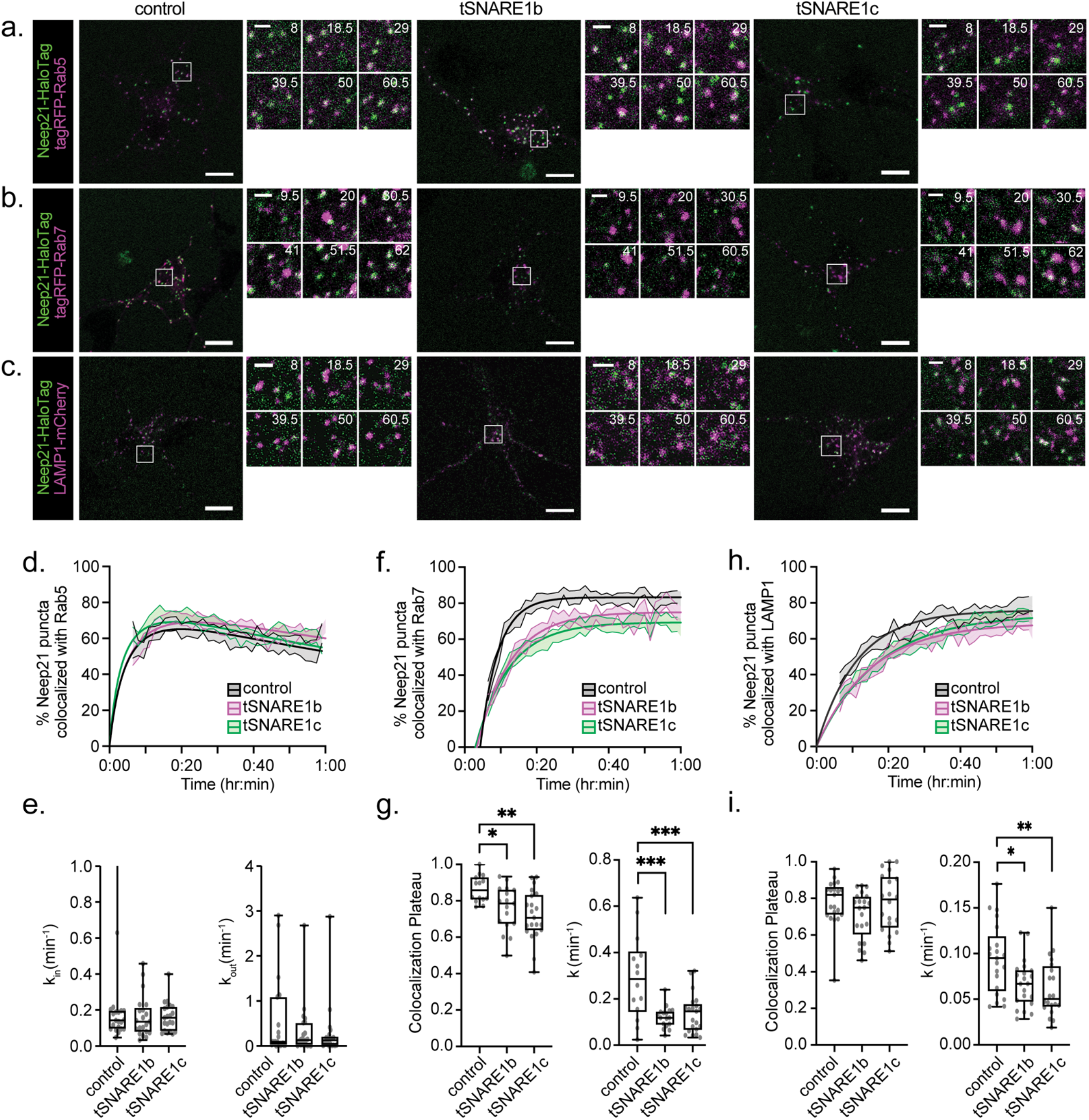
Overexpression of tSNARE1 delays trafficking of Neep21 into late endosomal and lysosomal compartments. (a-c) Representative single frames from movies of neurons at 2 *DIV* expressing Neep21-HaloTag and either (a) tagRFP-Rab5, (b) tagRFP-Rab7, or (c) LAMP1-mCherry, with or without the GFP-tSNARE1 isoforms listed. Scale bar: 10 μm (large), 2 μm (zoom montage) with time in minutes. (d) The percentage of Neep21 puncta colocalized with Rab5. n = 22 (control), 21 (tSNARE1b), 21 (tSNARE1c). (e) Neep21 entry into (k_in_) or exit out of (k_out_) Rab5^+^ compartments. (f) The percentage of Neep21 puncta colocalized with Rab7. n = 14 (control), 17 (tSNARE1b), 21 (tSNARE1c). (g) The colocalization plateau and rate constant for Neep21 entry into Rab7^+^ compartments. (h) The percentage of Neep21 puncta colocalized with LAMP1. n = 21 (control), 21 (tSNARE1b), 20 (tSNARE1c). (i) The colocalization plateau and the rate constant for Neep21 entry into LAMP1^+^ compartments. Data is reported as the mean curve fit model +/− SEM at each time point. For the quantification of rates and colocalization plateaus, data is reported as box and whiskers plot, with the min and max values as whiskers.

Neep21 rapidly colocalized with Rab5^+^ compartments and slowly left over an hour time period. There was no difference between the rate of Neep21 puncta entering or exiting Rab5^+^ early endosomes with or without expression of tSNARE1b or tSNARE1c (Fig. 6d&e, k_in_ p=0.3905, k_out_ p=0.4204). Strikingly, expression of either tSNARE1b or tSNARE1c delayed entry of Neep21 into Rab7^+^ compartments, and Neep21 never reached the extent of colocalization with Rab7 as it did in the absence of tSNARE1 (Fig. 6f&g, plateau p=0.0291 (tSNARE1b) & p=0.0016 (tSNARE1c), k p=0.0002 (tSNARE1b) & p=0.0004 (tSNARE1c)). Overexpression of tSNARE1b or tSNARE1c also delayed entry of Neep21 into LAMP1^+^ associated compartments (Fig. 6h&i, plateau p=0.1940 (tSNARE1b) & p=0.9960 (tSNARE1c), k p=0.0192 (tSNARE1b) & p=0.0051 (tSNARE1c)). As the only difference between tSNARE1b and tSNARE1c is the exclusion or inclusion of the Myb-like domain, this finding suggested that the Myb-like domain does not regulate Neep21 trafficking through the endosomal system. Therefore, the SNARE domain of tSNARE1 delayed trafficking of Neep21 into Rab7^+^ late endosomal and LAMP1^+^ lysosomal compartments.

## Discussion

Although many genetic loci have been implicated in schizophrenia, without understanding the function of the implicated gene products, the etiology of schizophrenia will remain unclear. *TSNARE1* is a high-confidence gene candidate for schizophrenia risk, but its biological function in the brain was previously unknown. We identified the four most abundant mRNA products of *TSNARE1*; the sequence information from which gave clues as to their likely biological function. Collectively, the *TSNARE1* mRNAs encoded two potential functional domains: a predicted syntaxin-like Qa SNARE domain, present in all four isoforms, and a Myb-like domain that is present in three of the four isoforms. Whereas the data we presented does not eliminate a possible role for the Myb-like domain in schizophrenia pathogenesis associated with *TSNARE1*, the weight of data presented here point most strongly to a role for tSNARE1 as a regulator of membrane trafficking in the endolysosomal system. This function is most likely key to both its normal activity within neurons as well as its dysfunction, which is associated with schizophrenia.

What are the features related to human *TSNARE1* that suggest that its biological function is linked to endocytic trafficking? First, all four major isoforms contained the Qa SNARE domain while only a subset of these included the Myb-domain, and it is the collective overexpression of these isoforms that is associated with schizophrenia in humans. Second, just as overexpression of tSNARE1 is associated with schizophrenia, we found that overexpression of either of the two most common forms of tSNARE1 in mouse primary neurons induced defects in endocytic trafficking. Third, recombinant forms of tSNARE1 could replace Stx12 in early endosome SNARE complexes—supporting a possible role for tSNARE1 as an inhibitor or i-SNARE (see below) for endocytosis in early endosome maturation into late endosomes. Finally, another endocytic regulator in neurons, *SNAP91*, was recently identified as a high confidence schizophrenia risk allele and appears to genetically interact with *TSNARE1*^2,30^. Taken together, these observations provide a compelling case for the role of tSNARE1 in schizophrenia being at the level of endocytic trafficking—likely in the endocytic trafficking or maturation of early endosomes to the late endosome.

Why then does the *TSNARE1* gene contain exons encoding a Myb-like domain? While we can not exclude a function for tSNARE1 in the nucleus, one likely explanation is that the *TSNARE1* gene arose during vertebrate evolution as a product of a Harbinger transposition^4,5^. Since the Myb-domain descended from the Harbinger transposon that is required for the transposition event itself, its presence may simply be a relic of this transposition. Further, because the Myb-like domain has several nuclear localization signals associated with the DNA-binding activity of this domain, it is possible that rather than functioning in the nucleus, the nuclear localization may act to regulate the partitioning of tSNARE1 in and out of the nucleus to control the level of tSNARE1 in the cytoplasm.

A majority (~93-97%) of tSNARE1 in the brain (three of the four isoforms) lacked a transmembrane domain or a predicted lipid attachment site, which is critical for membrane fusion facilitated by Qa-family SNARE proteins^31^. This is an important difference, as Qa SNAREs lacking a transmembrane domain would likely inhibit, rather than promote, membrane fusion. Therefore, tSNARE1 has the properties previously predicted by Rothman and colleagues of an i-SNARE (or inhibitory SNARE)^32^. i-SNAREs were proposed as a mechanism by which SNARE family proteins would act to modulate membrane trafficking events. This proposed mechanism would involve i-SNAREs negatively regulating trafficking by forming fusion-incompetent complexes, thereby acting as competitive inhibitors to the formation of canonical fusion-competent SNARE complexes. In the case of tSNARE1, the most likely target of this inhibition is Stx12, which is thought to function in endosomal trafficking. Our biochemical demonstration that tSNARE1c can indeed form ternary SNARE complexes with the same three endosomal SNAREs as Stx12 fits nicely with this possibility. Whereas i-SNAREs have been predicted and demonstrated *in vitro,* we believe this is likely the first example of an i-SNARE implicated in human disease. However, although our results are consistent with the hypothesis that tSNARE1 acts as a negative regulator to membrane fusion, studies in a reconstituted system such as liposomal fusion assays will be important to directly test tSNARE1 as an i-SNARE.

Previous reports suggested tSNARE1 is likely to function in synaptic vesicle exocytosis^30,33–35^. However, this statement is most likely related to confusion over classification of tSNARE1 as a “target” SNARE protein based on it being related to the syntaxin family Q-SNARE proteins, whereas its specific role in the cell was completely uninvestigated. Whereas the syntaxin family member Stx1 does indeed have a well-documented role in synaptic vesicle exocytosis, the sequence of tSNARE1 is most closely associated with Stx12, which is thought to be involved in endosome:endosome fusion necessary for maturation of early endosomes into late endosomes^36^. Therefore, tSNARE1 is highly unlikely to function in synaptic release directly. Indeed, we showed that tSNARE1 is localized to the endolysosomal system rather than to the plasma membrane—where Stx1 is found^37^. Further, these previous studies use, to our knowledge, a human isoform of tSNARE1 which contains a TM domain (tSNARE1a), which we find only represents 3-7% of tSNARE1 in human brain.

How tSNARE1 dysfunction contributes to schizophrenia pathogenesis is not completely understood. Patients with schizophrenia have less dendritic spines than neurotypical patients^24^, and lysosomes have recently been shown to traffic to dendritic spines in an activity-dependent manner^38^. tSNARE1 may therefore be poised to function in the post-synapse, perhaps to regulate receptor availability or protein homeostasis. tSNARE1 may also function at the pre-synapse, perhaps by altering the availability of machinery needed for synaptic release. The fact that overexpression of tSNARE1 is linked to schizophrenia suggests that the membrane trafficking events that tSNARE1 regulates require precise levels of tSNARE1, whose dysfunction likely imparts changes at the synapse. Determining the specific trafficking events that tSNARE1 regulates and how tSNARE1 imparts changes at the synapse may reveal common pathways disrupted in schizophrenia and possible targets for drug design and clinical intervention.

## Materials and Methods

### Animals

Wild-type C57BL/6J background mice were purchased from The Jackson Laboratory and bred at the University of North Carolina at Chapel Hill (UNC). Mice are genotyped to confirm wild-type identity with standard genotyping procedures. Mice are kept pathogen-free environments with 12-h light and dark cycles with ready access to food and water. Male and female mice were placed in a cage together overnight and timed pregnant females were identified at E0.5 if the female had a vaginal plug. All procedures and husbandry were conducted according to protocols approved by the Institutional Animal Care and Use Committee at UNC.

### Plasmids

Stx6 human cDNA (BC009944), residues 1-255 in pGEX6P1 (pB2247). Stx12 human cDNA (BC046999), residues 1-278 in pGEX6P1 (pB2249). VAMP4 human cDNA (BC005974), residues 1-141 in pGEX6P1 (pB2258). Vit1a human cDNA (NM145206), residues 1-217 in pET15b (pB2246). Synthetic tSNARE1 (NP 001353833.1 codons optimized for *E.coli* expression), residues 1-496 in pGEX4P1. tSNARE1a human cDNA (NM_145003.5) in pEGFP-C2 (pB2468). tSNARE1b human cDNA (NM_001291931.1) in pEGFP-C2 (pB2466). tSNARE1c human cDNA (NM_001366904.1) in pEGFP-C2 (pB2467). Stx12 human cDNA (BC046999) in pmCherry-C1 (pB2265). Human Rab7 in mTag-RFP-N1 (pB2357). Mouse Neep21 in Halo-N1 (pB2504), cloned from Neep21-mCherry, a gift from B. Winckler (University of Virginia). TagRFP-Rab4a, TagRFP-Rab5a, TagRFP-Rab11a were gifts from J.S. Bonifacino (NIH). LAMP1-mCherry and Halo-N1 were a gift from J. Lippincott-Schwartz (HHMI Janelia). All unique materials are readily available from the corresponding author upon request.

### Antibodies

Antibodies against: anti-tSNARE1 was from LifeSpan Biosciences, Inc. (cat # LS-C160258) rabbit polyclonal Ab used at 1:1000 dilution for immunoblot analysis; anti-Stx6 was from Cell Signaling Technology (cat #28696) rabbit monoclonal Ab C34B2 used at 1:1000 dilution for immunoblot analysis; anti-Stx12 LifeSpan Biosciences, Inc. (cat # LS-B2989) mouse monoclonal Ab used at 1:1000 dilution for immunoblot analysis.

### *TSNARE1* mRNA Identification

Human Brain Total RNA (Takara Cat. #636530) and Human Brain Cerebral Cortex Total RNA (Takara Cat. #636561) was reverse-transcribed by Maxima H Minus First Strand cDNA Synthesis Kit (Thermo Scientific) with oligo (dT)_18_ primers (1 μg RNA for each 20 ul reaction). The primers (Eurofins) used for *TSNARE1* transcript amplification are as follows: TSN1-NT3 (5’ - gcatcaggatccaaatcgcccgtggaggtggc – 3’), TSN1-CT3 (5’ – gcatcagtctagatcaggggcagccacaagcagctgggggt - 3’), TSN1-CT2b (5’ - gcatcagtctagatcactttcggacagaggtggcgatgatgatga - 3’). 1 μL cDNA was amplified using 1 μL Phusion High-Fidelity DNA Polymerase (NEB) in HF Phusion buffer with 10 μM primers, 200 μM dNTPs (Agilent), +/− 5% DMSO per 50 μl reaction. Transcripts were checked on agarose gels, purified with phenol:chloroform:isoamyl alcohol (PCI) and Wizard resin (Promega), digested with XbaI and BamHI-HF in Cutsmart Buffer (NEB), and gel purified using Gel Extraction Kit (Qiagen). Isolated transcripts were ligated into pEGFP-C2 using Quick Ligation Kit (NEB), cloned into DH5α, purified with Wizard Genomic DNA Purification kit (Promega) and confirmed via sequencing (GeneWiz). tSNARE1a (NM_145003.5), tSNARE1b (NM_001291931.1), and tSNARE1c (NM_001366904.1) matched predicted isoforms on Refseq. tSNARE1d did not.

### Transcriptomic Data Analysis

STAR aligned bam files of adult prefrontal cortex RNA-sequencing data was downloaded from PsychENCODE^2^ and visualized with Integrative Genomics Viewer (IGV) Viewer (Broad Institute). Sashimi plots were modified from IGVViewer (Broad Institute) to display exon coverage and the number of reads connecting two exons. Fetal cortical wall bam files were generously supplied from Dr. Jason Stein^19^ (UNC) and analyzed as above. To determine the expression of *TSNARE1* in different cell types, we utilized single-cell transcriptomic data from the adult brain^39,40^.

### Protein Purification and SNARE Assembly

Recombinant human Syntaxin6, Syntaxin12 and tSNARE1 were purified as previously described^41^ except the GST-fusion proteins were cleaved with HRC 3C protease (Pierce) overnight at 4C in buffer containing 50mM Tris pH 7.5, 150mM NaCl, 1mM EDTA, 0.5% Triton X-100 and 0.5mM DTT. Recombinant human Vti1a was purified using His select nickel affinity column following manufactures’ directions and dialyzed into buffer containing 20mM Tris pH 7.5, 150mM NaCl, 10% glycerol and 0.5% Triton X-100. Various combinations of soluble recombinant SNARE proteins (Syntaxin6, Vti1a, and Syntaxin12 or t-SNARE1) were mixed with GST-Vamp4 immobilized on glutathione Sepharose beads in 100μl reactions with buffer containing 50mM Tris pH 7.5, 150mM NaCl, 3mMMgCl2, 0.5% Triton X-100 and 0.5mM DTT. All soluble proteins were present at 1μM final concentration while the GST-Vamp4 beads were present at 0.6μM final concentration. Soluble proteins were preincubated on ice for 20 minutes with Sepharose beads and then centrifuged at 13,000 rpm for 15 minutes at 4C before adding to GST-Vamp4. The binding reaction was incubated overnight at 4C and the supernatants were then separated from the pellets by centrifugation. The pellets were washed four times in binding buffer and then boiled in SDS sample buffer for analysis by Coomassie and immunoblot.

For pull down of heterologous SNARE complexes, mouse E15.5 embryo brains were dissected and lysed in 20mM Tris pH 7.5, 140mM NaCl, 5mM MgCl2 5% glycerol and 0.5% Triton using a dounce homogenizer. Lysate was left on ice 20 minutes and then spun at 13000 rpm at 4C for 15 minutes. Lysate concentration was determined^42^, and adjusted to 25mg/ml and before adding to immobilized beads of GST, GST-Vamp4 or GST-tSNARE1 at a final concentration of 1.5 μM. Binding was conducted at 4C for 2 hours and then the supernatant was separated from the beads and the beads washed four times in binding buffer before boiling in sample buffer and analyzing by western blot analysis.

### Cortical Neuron Culture

Cortices were microdissected from both male and female E15.5 mouse embryos and either were plated immediately or stored in Hibernate E (ThermoFisher) and plated within 24 hours. Cortical neurons were dissociated with trypsin and plated on poly-D-lysine (Sigma)-coated imaging dishes (MatTek for colocalization, 8-well CellVis cover glass for Neep21 trafficking) and grown and imaged in serum-free Neurobasal medium (Gibco) supplemented with B27 (Invitrogen) and L-glutamine. To transfect, neurons are centrifuged and resuspended in transfection reagent (Amaxa Nucleofector; Lonza) and a nucleofector is used to electroporate the neurons according to the manufacturer protocol prior to plating on imaging dishes. To quantify colocalization of tSNARE1 with endosomal markers, 1 μg of each GFP-tSNARE1 isoform was cotransfected with 0.5-1 μg of spectrally distinct red-tagged markers of the endosomal system (tagRFP-Rab4, tagRFP-Rab5, tagRFP-Rab11, tagRFP-Rab7, LAMP1-mCherry, and mCherry-Stx12). The Neep21 trafficking experiments were performed by transfecting 1μg Neep21-HaloTag and 1μg of either tagRFP-Rab5, tagRFP-Rab7, or LAMP1-mcherry, with or without 1μg GFP-tSNARE1. Directly prior to imaging, 1 μL 3.5 mM Alexa Fluor 660 HaloTag ligand (Promega) was added to 200-300 μL cells and incubated at 30°C for 5 minutes. The neurons were washed twice with fresh medium and imaged.

### Microscopy

Pyramidal cortical neurons were imaged at two days *in vitro* by live-cell confocal microscopy with an inverted 880 laser scanning microscope (Zeiss) with a 32-channel GaAsP spectral detector and a Plan Apo 63x 1.4 NA oil objective (Zeiss) in a humidified chamber at 37°C and 5% CO_2_. Simultaneous, bidirectional scans of z-stacks with dual or triple channels, tight gate settings, and a 1-AU pinhole were acquired using the Zeiss ZEN software. For Neep21 trafficking assays, the chamber was set to 30°C and the pinhole was increased to maximize detection of endosomal surfaces. Movies of multiple neurons were recorded simultaneously, using the Positions function on the ZEN software and imaging once every 90 seconds for an hour.

### Image Analysis

For colocalization analysis, a z-slice containing the majority of tSNARE1 puncta was chosen for quantification. Custom image-analysis software developed by the Taraska^27^ lab was used to measure colocalization of puncta using MATLAB (MathWorks). The program generates a region of interest (ROIs) of 7 pixels radius around tSNARE1 (green) puncta and then uses the same ROIs in the marker (red) channel. It computes the colocalization in these ROI based off two metrics, correlation (C), or the average Pearson’s values of the ROIs, and peak height (h), which is the maximum intensity difference between the central pixel in the other (red) channel of the ROI and any other pixel in that ROI. Perfect colocalization would have a C and h value of 1.0, whereas perfect exclusion would have a C and h value of −1.0. For Neep21-HaloTag analysis, Imaris (Oxford Instruments) was used to quantify the percentage of Neep21 puncta coincident with an endosomal marker over time. Movies were included if they were started by 9 minutes and 30 seconds post dye addition, did not bleach before the end of the movie, and had good signal:noise. For Neep21, we automatically detected puncta over time using the Spots feature and manually removed points if picked up from fluorescence outside the cell. We used the Surfaces feature to automatically detect endosomal surfaces (tagRFP-Rab5, tagRFP-Rab7, or LAMP1-mCherry). We counted a Neep21 puncta within an endosomal surface as long as its center point was within.03 μm from an endosomal surface. From this, we calculated the percentage of Neep21 within an endosomal compartment for each time point. We rounded each time point to the nearest minute and a half increment in order to graph and analyze the data using GraphPad Prism.

### Conservation Analysis

We compared the phyloP scores between each domain, which measures nucleotide substitution rates that are more reduced (conservation) or increased (acceleration) than expected under neutral drift^43^. We compared the core SNARE domain (hg38 “-” strand, chr8:142300633-142300647, chr8:142284413-142284485, chr8:142274781-142274851) with the Myb-like domain (hg38 “-” strand, chr8:142344079-142344414). Because *TSNARE1* arose early in vertebrate evolution, we compared the phyloP scores for each of these domains using the 100 vertebrate pairwise alignment Conservation track from UCSC Genome Browser and reported as the mean +/− SEM. Minor allele frequency was downloaded through the Ensembl genome browser and the locations of the SNARE and Myb-like domain (above) were compared. Copy number variation (CNV) was reported using data from the Exome Aggregation Consortium (ExAC) for each of the exons of *TSNARE1*. For the protein alignment between tSNARE1 and Stx12, we used DNASTAR (DNASTAR).

### Statistics and Visualization

Multiple biological replicates were performed, and for imaging experiments multiple neurons were images within each biological replicate. Graphs were either generated with Microsoft Excel, Past4, or GraphPad Prism software and modified with Adobe Illustrator. ImageJ software^44–46^ (National Institutes of Health) was used for image visualization. Power analysis was performed for colocalization analysis using preliminary colocalization data. Experimental condition was blinded for analysis. For Neep21 trafficking into Rab7^+^ or LAMP1^+^ compartments, the data were fit to a non-linear regression one phase association model. For Neep21 trafficking into Rab5^+^ compartments, a one phase association model was separately fit to the data before (k_in_) or after (k_out_) the maximum percentage of Neep21 in Rab5^+^ compartments. The k and plateau values from these curve fits were plotted and compared. We assumed normal Gaussian distributions for all data. We compared groups using an ordinary one-way ANOVA with Dunnett’s test for multiple comparisons against the control with 95% confidence intervals. The exact p values are listed in the text. For Neep21 with Rab5, F (DFn, DFd) = 0.9551 (2, 61) for k_in_ and F (DFn, DFd) = 0.8787 (2, 62) for k_out_. For Neep21 with Rab7, F (DFn, DFd) = 11 (2, 49) for k and F (DFn, DFd) = 6.526 (2, 49) for plateau. For Neep21 with LAMP1, F (DFn, DFd) = 5.729 (2, 59) for k and F (DFn, DFd) = 1.652 (2, 59) for plateau.

## Acknowledgements

This work is supported by grants from National Institutes of Health grants R01GM054712-23 to P.B., R01NS105614 and R35GM135160 to S.L.G,, R00MH113823 and DP2MH122403 to H.W., and F31MH116576 and T32GM119999 to M.P. We would like to thank Dr. Patrick Sullivan for contributions during the early phases of this work. We are indebted to PsychENCODE, Dr. Jason Stein, Dan Liang, Mike Lafferty, and Joel Parker of the Lineberger Bioinformatics Core for their assistance in quantifying RNA-sequencing data. We thank Dr. Daniel Schrider for his advice on comparing the conservation of tSNARE1 domains. We would like to thank the Exome Aggregation Consortium and the groups that provided exome variant data for comparison. The full list of contributing groups can be found at http://exac.broadinstitute.org/about.The Genotype-Tissue Expression (GTEx) Project was supported by the Common Fund of the Office of the Director of the National Institutes of Health, and by NCI, NHGRI, NHLBI, NIDA, NIMH, and NINDS. The data used for the analyses described in this manuscript were obtained from the GTEx Portal. We thank Robert Currin the Hooker Imaging Core at UNC.

## Data Availability

The datasets generated during and/or analyzed during the current study are available from the corresponding authors on reasonable request.

---

**Supplementary Fig. 1:**
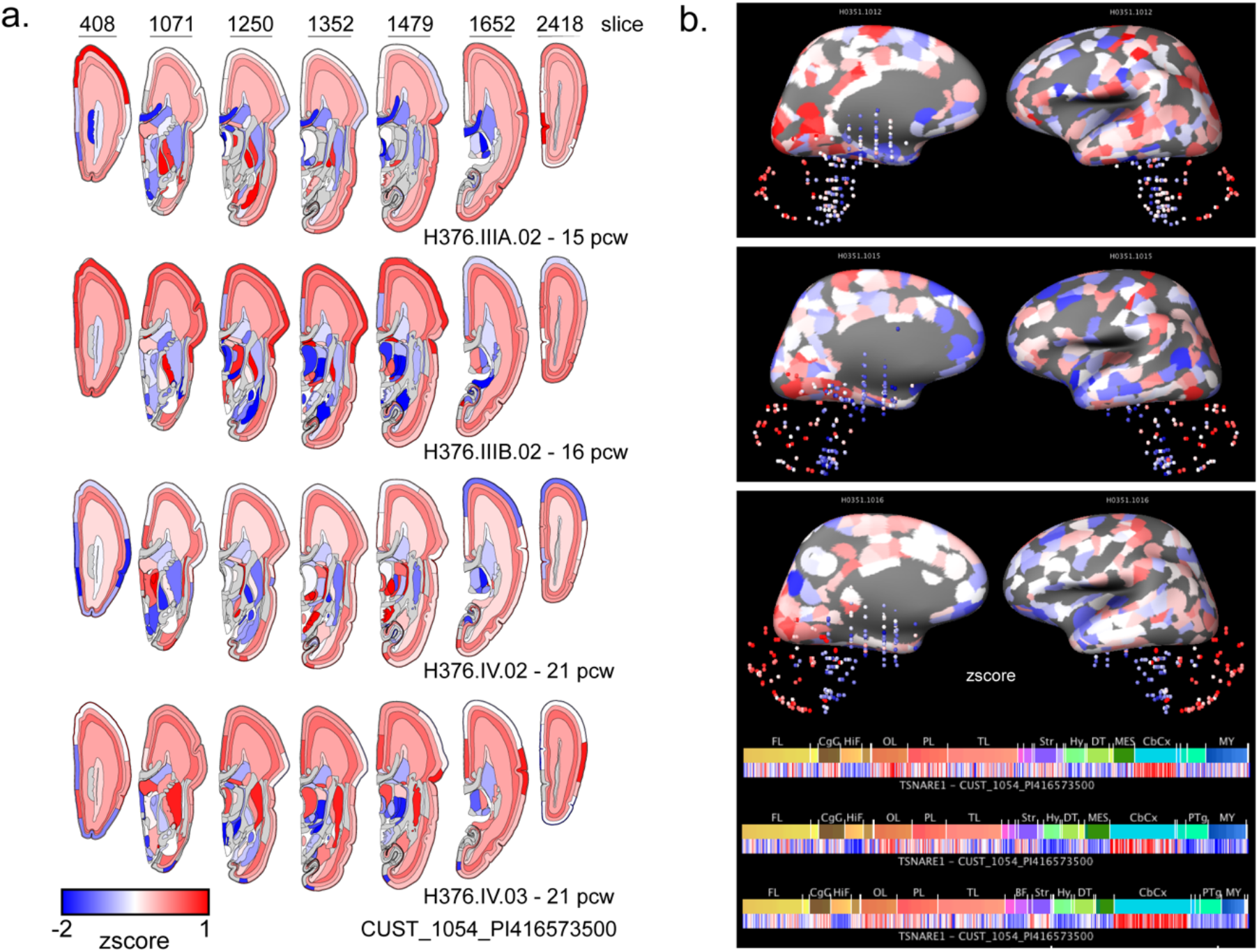
*TSNARE1* gene expression by brain region. Brain microarray data displayed using the visualization tools on Allen Brain Atlas for tSNARE1 enrichment in both (a) fetal and (b) adult brain.

**Supplementary Fig. 2:**
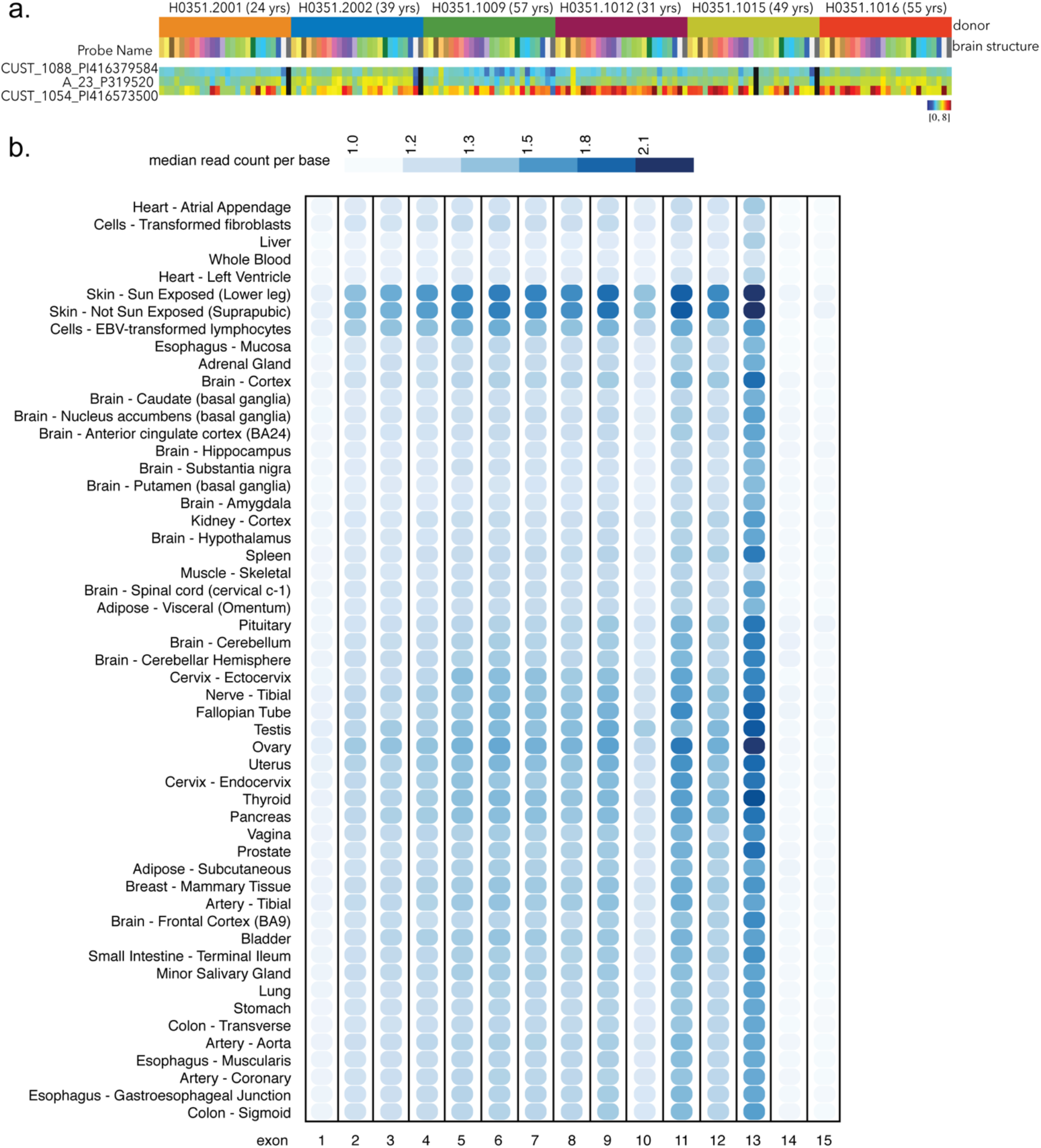
*TSNARE1* is alternatively spliced and the majority of transcripts lack a TM domain. (a) Heat map of the log2 intensity of the probe common to all *TSNARE1* isoforms (CUST_1054_P1416573500) and those probes that are contained within exon 14 (CUST_1088_P1416379584, A_23_P319520). (b) Heat map of the median read count per base from data adapted from GTEx.

**Supplementary Fig. 3:**
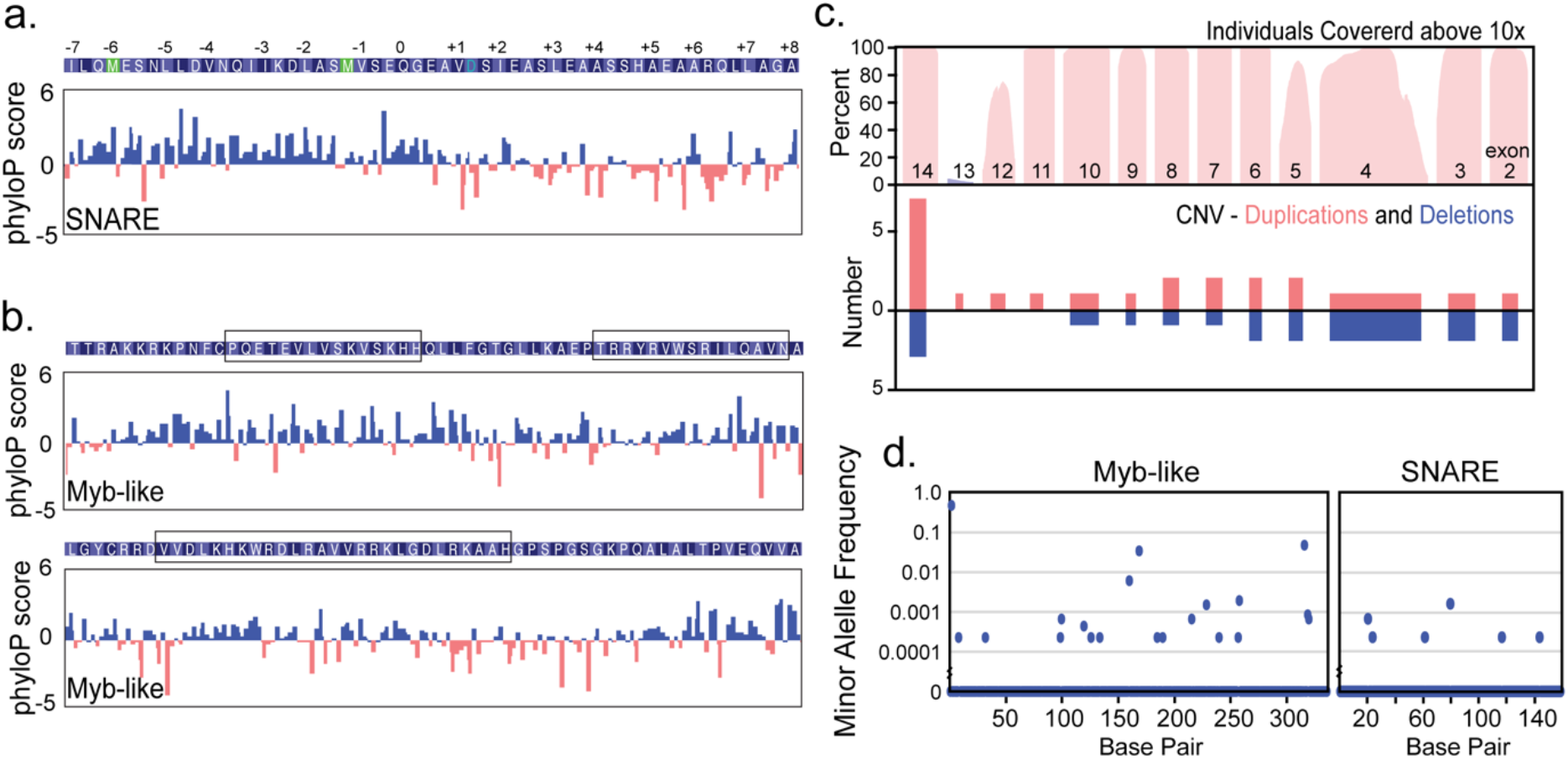
The SNARE and Myb-like domains of tSNARE1 are conserved. (a&b) phyloP scores computed from 100 vertebrate alignment per base of the (a) SNARE domain with the layers of interacting amino acids within the four-helical bundle marked and (b) the Myb-like domain with the predicted three helices boxed. (c) The percent of individuals with sequencing coverage above 10x at each base (top panel) and copy number variation (CNV) duplications and deletions (bottom) from the ExAC database. (d) The minor allele frequency (MAF) per base of each domain.

**Supplementary Fig. 4:**
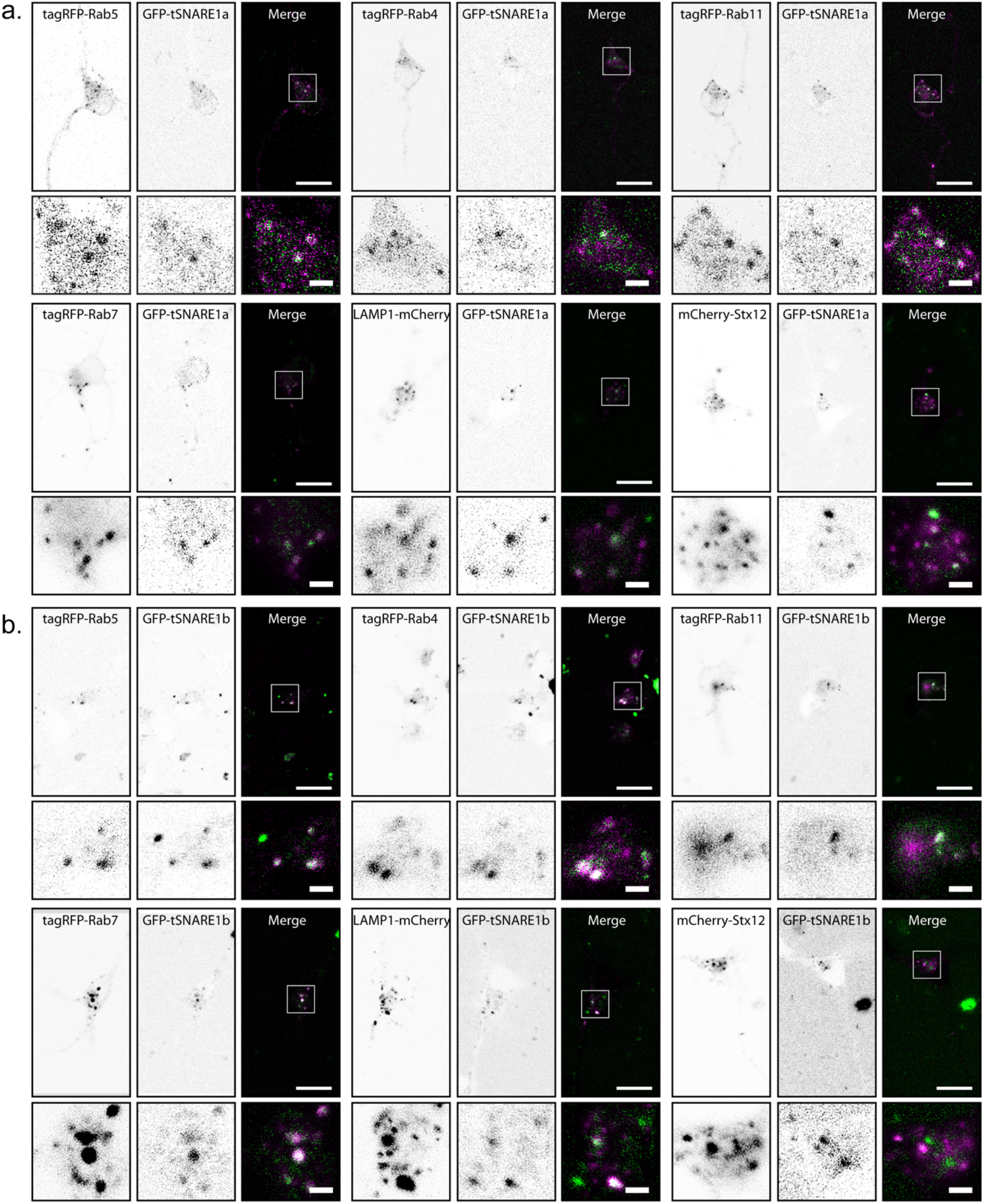
tSNARE1a and tSNARE1b localize to the endosomal network. (a-b) Example images of murine cortical neurons transfected with (a) GFP-tSNARE1a or (b) GFP-tSNARE1b and various red-tagged endosomal markers, with a zoomed in region of the soma marked by a white box. Scale bar=15μm, 2.5μm (zoom).

**Movie 1: Neep21 colocalization with Rab5 over time.**

**Movie 2: Neep21 colocalization with Rab7 over time.**

**Movie 3: Neep21 colocalization with LAMP1 over time.**

